## Supplementary figures for "A kidney-hypothalamus axis promotes compensatory glucose production in response to glycosuria"

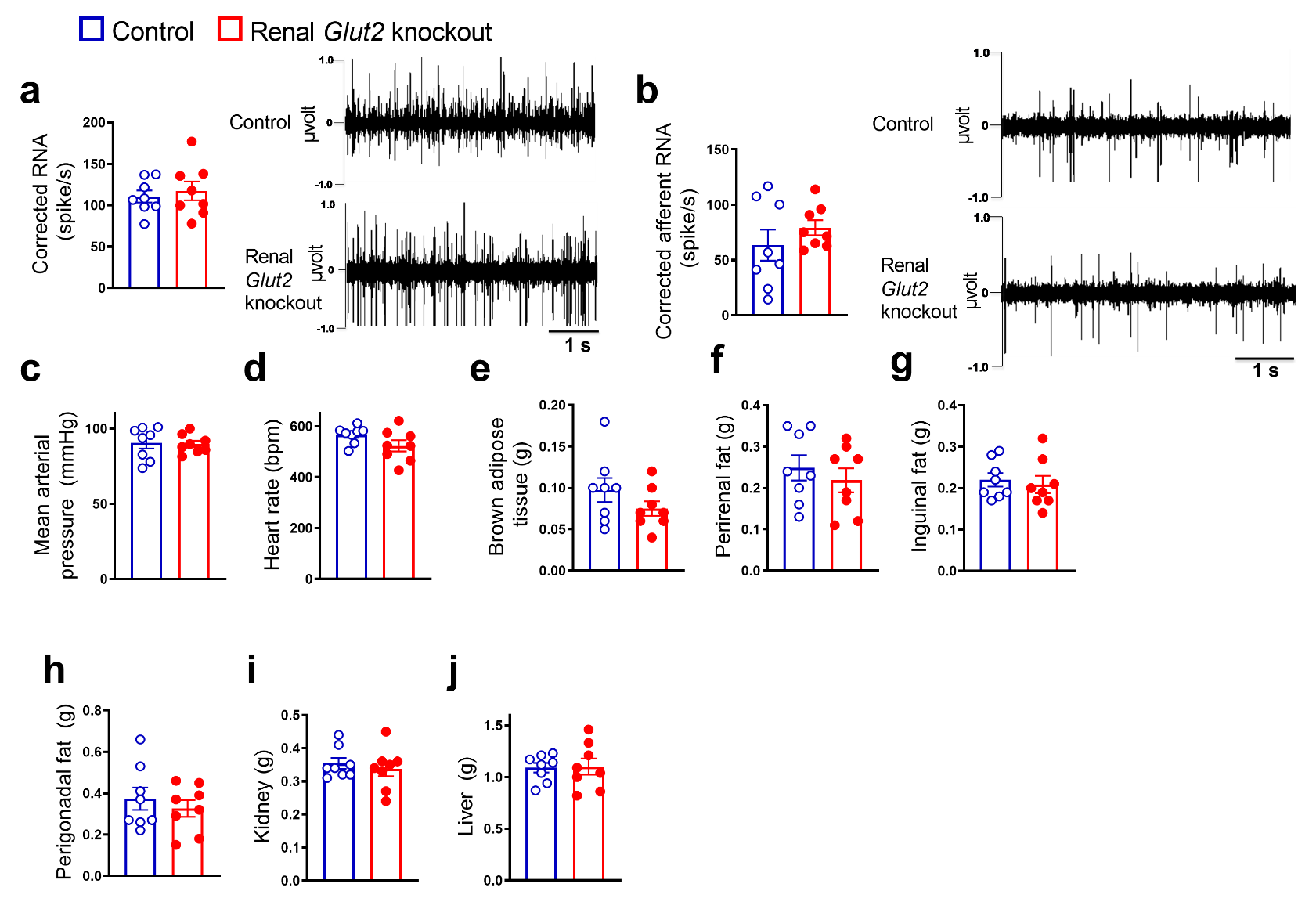

**Supplementary figure 1:** Total and afferent renal nerve activity including the representative traces **(a,b)**, mean arterial pressure and heart rate **(c,d)**, weights of brown **(e)** and regional white **(f-h)** adipose tissues, kidney **(i)**, and liver **(j)**, in 30 weeks old female renal *Glut2* knockout and their littermate control mice 16 weeks after inducing the *Glut2* deficiency.

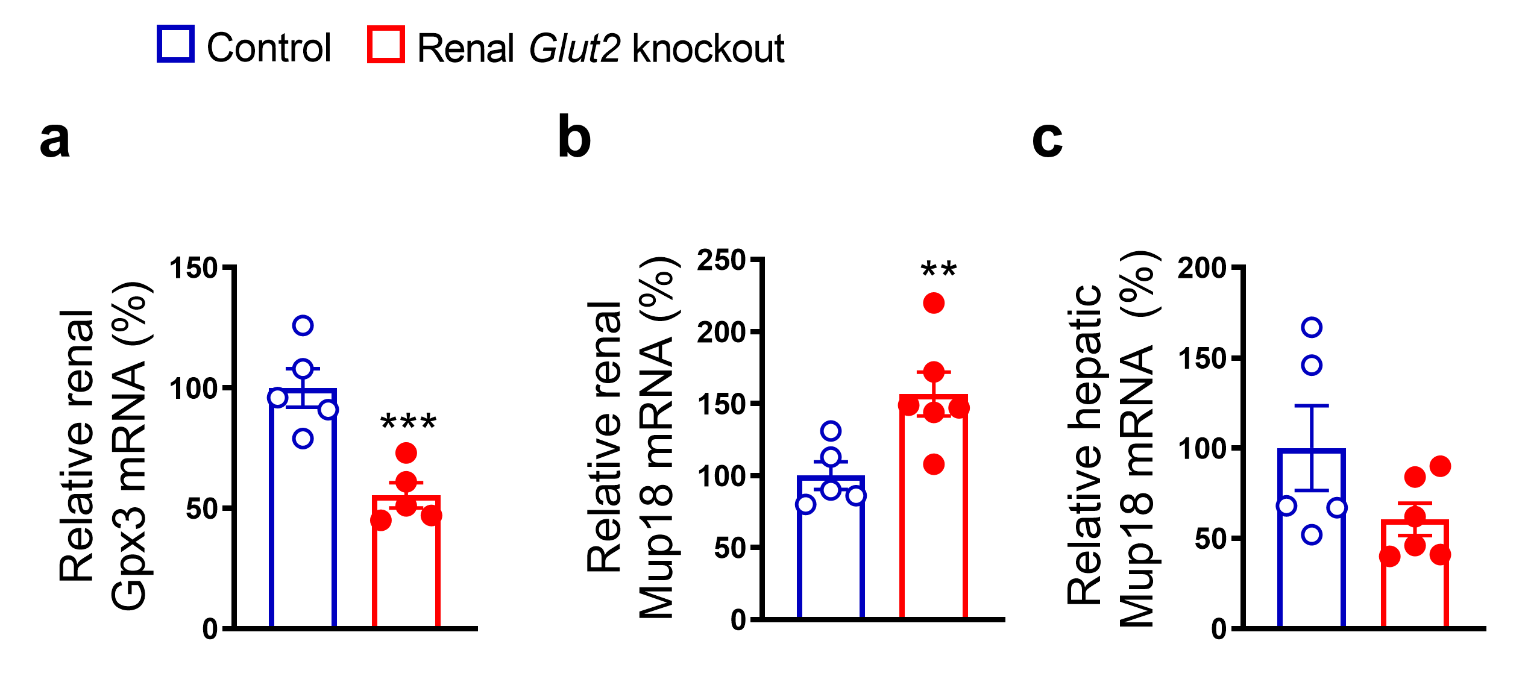
**Supplementary figure 2:** Renal *Glut2* knockout male mice (28 weeks old) have reduced expression of glutathione peroxidase 3 (Gpx3) **(a)** and major urinary protein 18 (Mup18) **(b)** in the kidneys without affecting hepatic Mup18 **(c),** measured by RT-qPCR 12 weeks after inducing the *Glut2* deficiency. **p<0.01, ***p<0.001, unpaired 2-tailed Student’s t-test. Data are presented as mean ±sem.
